# Bayesian mapping reveals that attention boosts neural responses to predicted and unpredicted stimuli

**DOI:** 10.1101/084517

**Authors:** M. I. Garrido, E. G. Rowe, Veronika Halász, J. B. Mattingley

**Author notes:** Correspondence: Centre for Advanced Imaging, The University of Queensland Building 57, Research Road, St Lucia 4072, Brisbane, Australia.

## Abstract

Predictive coding posits that the human brain continually monitors the environment for regularities and detects inconsistencies. It is unclear, however, what effect attention has on expectation processes, as there have been relatively few studies and the results of these have yielded contradictory findings. Here, we employed Bayesian model comparison to adjudicate between two alternative computational models. The *Opposition* model states that attention boosts neural responses equally to predicted and unpredicted stimuli, whereas the *Interaction* model assumes that attentional boosting of neural signals depends on the level of predictability. We designed a novel, audiospatial attention task that orthogonally manipulated attention and prediction by playing oddball sequences in either the attended or unattended ear. We observed sensory prediction error responses, with electroencephalography, across all attentional manipulations. Crucially, posterior probability maps revealed that, overall, the *Opposition* model better explained scalp and source data, suggesting that attention boosts responses to predicted and unpredicted stimuli equally. Furthermore, Dynamic Causal Modelling (DCM) showed that these *Opposition* effects were expressed in plastic changes within the mismatch negativity network. Our findings provide empirical evidence for a computational model of the opposing interplay of attention and expectation in the brain.

## INTRODUCTION

The way in which we perceive the world around us is thought to be an active inferential process. Rather than passively registering information that arrives at our senses, the brain builds predictive models of what it might encounter next. These theoretical conjectures have been formalised in terms of predictive coding (Rao RP and DH Ballard 1999; Friston K 2005) and are useful in explaining the ubiquitous phenomenon of larger brain responses to surprising than predictable events (Montague PR 1999; Garrido MI et al. 2013) (Opitz B et al. 1999; Summerfield C and E Koechlin 2008). Selective attention is the process of prioritising information by allocating more cognitive resources to the object of focus, while suppressing information that is irrelevant. Recent extensions of predictive coding have framed attention as the process of enhancing the reliability of prediction errors (Feldman H and KJ Friston 2010). This idea has been empirically demonstrated by larger prediction errors for attended than unattended visual objects (Jiang J et al. 2013) and sounds (Auksztulewicz R and K Friston 2015), with the latter going against the longstanding notion of mismatch negativity (MMN) as a pre-attentive process (Naatanen R et al. 2001).

There is a general consensus that expectation dampens neuronal activity and that attention boosts neuronal activity (Summerfield C and E Koechlin 2008). Thus, superficially at least, attention and prediction appear to have *opposing* effects. However, the way in which attention interacts with expectation is unclear for two reasons. First, there have been very few attempts to manipulate attention and prediction independently, but many instances in which the two have been entwined or confounded (Summerfield C and T Egner 2009), as attention is often manipulated in a probabilistic manner rather than through stimulus filtering or prioritisation. Second, the few studies on prediction and attention have yielded a puzzling depiction of what might be happening in the brain. Kok P et al. (2012) provided fMRI evidence that attention and prediction have an *interactive* or synergetic effect by showing greater brain activity in the visual cortex for predicted (than unpredicted) visual stimuli, a finding which was conceptually replicated using electroencephalography (EEG) for auditory stimuli, and expressed in the N1 evoked potential (Hsu YF et al. 2014). By contrast, (Auksztulewicz R and K Friston 2015) found that attention increased the typically observed difference between evoked responses to unpredicted versus predicted stimuli, as reflected in an enhanced MMN.

In this paper, we first formalise two theoretical models that have been put forward to explain the interplay between attention and prediction in the brain: the *Opposition* model and the *Interaction* model, introduced in Kok P *et al.* (2012). The *Interaction* model postulates that attention and prediction interact such that neuronal activity is greatest for attended and predicted events. This model is inspired by the idea that attention increases the precision of predictions by weighting prediction errors (Feldman H and KJ Friston 2010), and assumes four levels of precision, or attention, that depend on the level of prediction. By contrast, the *Opposition* model posits that attention and prediction have *opposing* effects on neural activity, such that prediction mitigates and attention boosts neural activity. The predictions of this model are that the neuronal responses will be greatest for attended unpredicted stimuli, and smallest for unattended predicted stimuli. Computationally, this model assumes that neuronal activity is weighted by two (instead of four) levels of attention (attended and unattended). This model is agnostic about the relationship between responses to attended predicted and unattended unpredicted events. Both the Interaction and the Opposition models assume that prediction has two levels, such that unpredicted stimuli evoke a larger neuronal response than predicted stimuli. They differ, however, in their treatment of the attention component. Specifically, the Opposition model offers a more parsimonious expression of the effects of attention on neuronal responses (see Figure 1).

**Figure 1.**
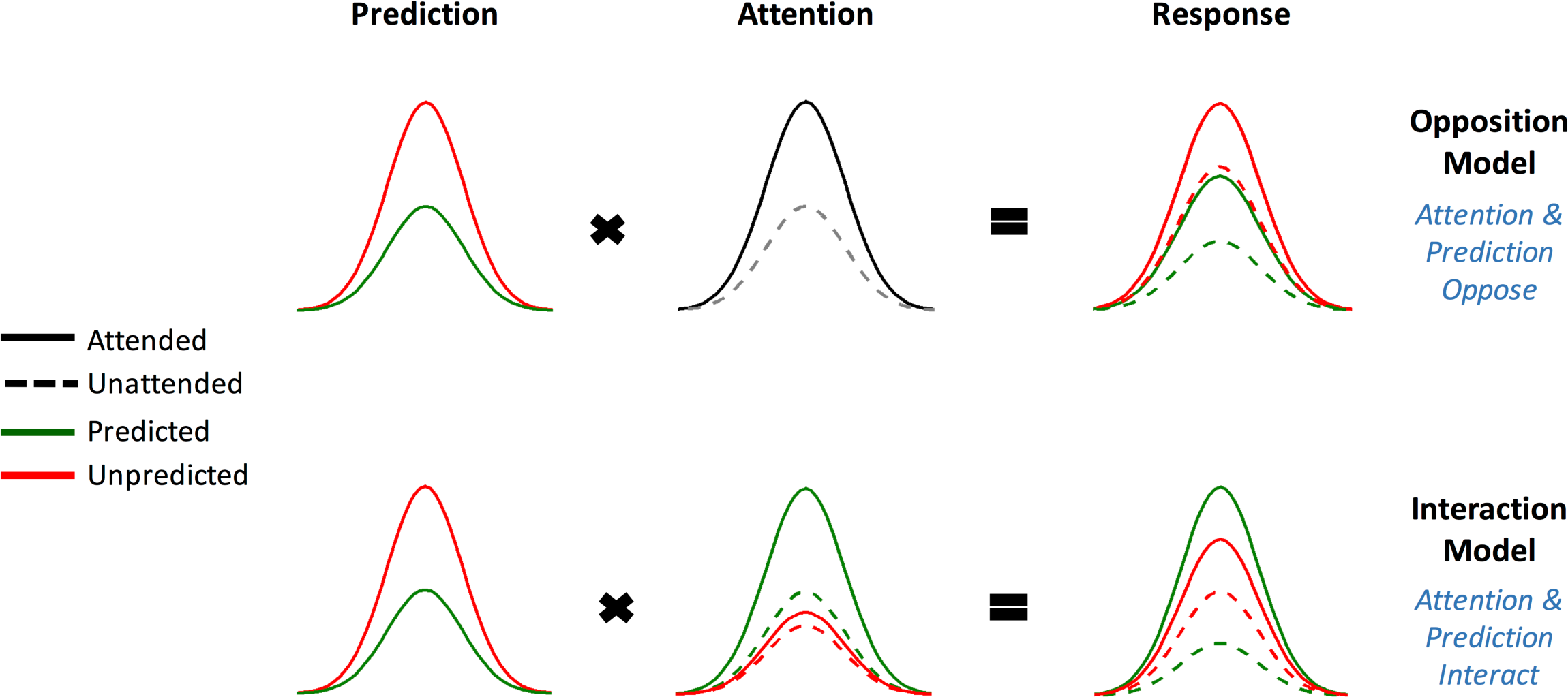
Two competing models for the relationship between attention and prediction. In the *Opposition* Model, predicted (green) and unpredicted (red) neural signals are multiplied by two levels of Attention, with attended stimuli (solid lines) receiving a greater boost than unattended stimuli (dashed lines). In the *Interaction* Model (proposed by Kok et al., 2011), predicted and unpredicted signals are multiplied by 4 (instead of 2) levels of Attention that depend on the level of the Prediction.

Here we tested these models empirically using Bayesian model comparison for scalp and source EEG data, as well as dynamic causal modelling (DCM). We developed a novel auditory task in which participants were presented with independent streams of white noise concurrently in each of the two ears, and were instructed to attend to the left channel, the right channel or both channels in separate blocks to detect brief gaps in the noise streams. At the same time, an irrelevant stream of standard and deviant tones was presented in either ear (attended or ignored), providing an orthogonal stimulus set from which to extract neural responses to predicted and unpredicted auditory events.

## METHODS

### Participants

Twenty one healthy adults were recruited for the experiment. Data from two participants were excluded from further analysis due to poor performace on the behavioural task (accuracy < 50%). The reported analysis was thus performed on data from 19 participants (10 females, aged 19-43, *M* = 24.21, *SD* = 6.11) with no reported history of neurological or psychiatric disorder and no previous head trauma resulting in unconsciousness. All participants gave written informed consent in accordance with the guidelines of the University of Queensland’s ethical committee, and were monetarily compensated for their time.

### Auditory Stimuli

The auditory task developed for the study is depicted in Figure 2. An auditory frequency oddball sequence was played to one ear at 60 dB and overlayed with Gaussian white noise at 40 dB. White noise only was played to the other ear at 40 dB. Two pure tones, standards (*p* = 0.85) and deviants (*p* = 0.15), (*f* = 500 or 550 Hz; counterbalanced between blocks) of 50 ms in duration were played with an inter-stimulus interval of 450 ms. Embedded in the white noise of either ear were two types of targets: a total of 30 non-overlapping randomised periods of no sound (*gaps*), which could be singular (90 ms gaps only, 15 per block) or doubled (two 90 ms breaks separated by a 30ms white noise return, 15 per block). The gaps in the white noise of either ear were never within 2.5 seconds of each other and never occurred at the same time as a tone. All auditory stimuli were created using in-house Matlab scripts, recorded using Audacity Sound Mixer prior to the experiment, and delivered with inner-ear buds (Etymotic, ER3).

**Figure 2.**
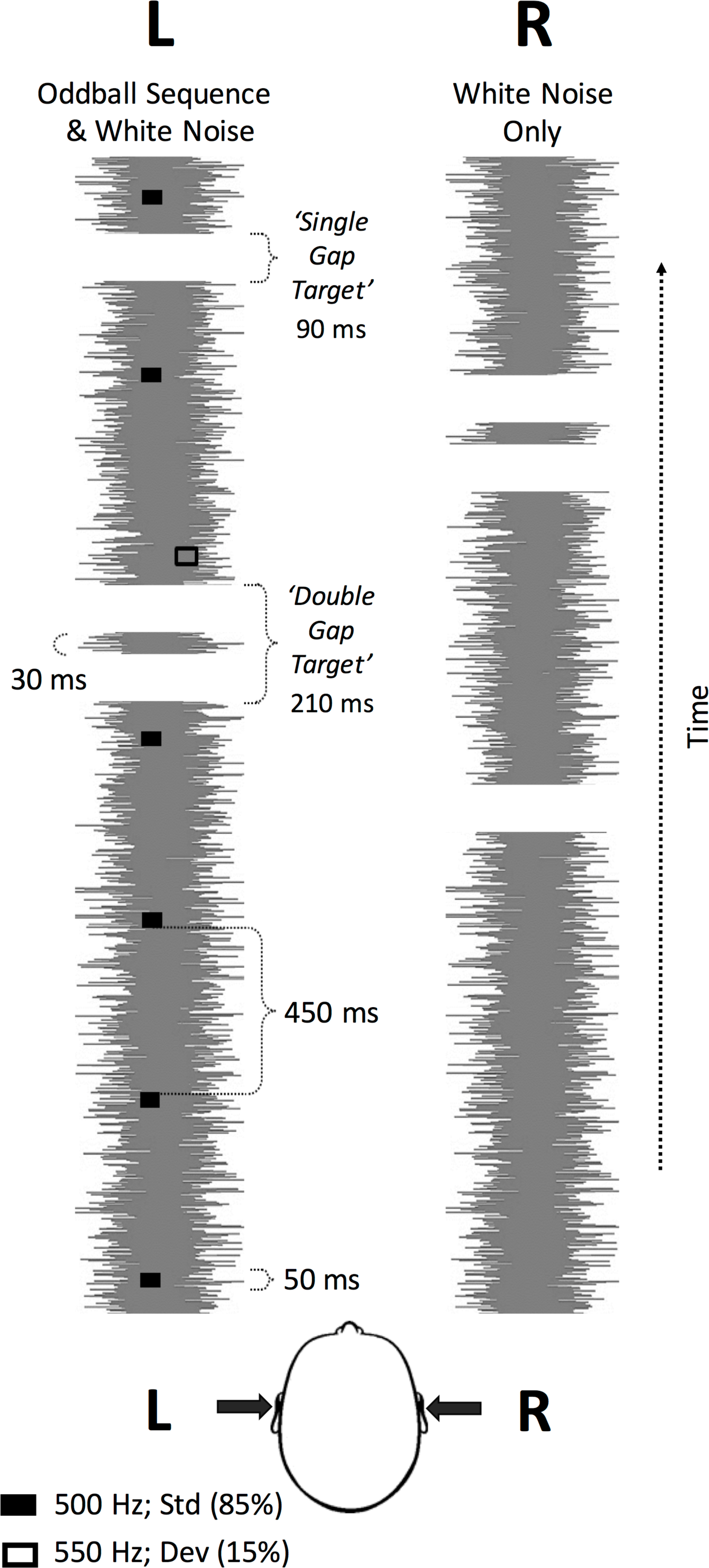
Experimental paradigm. Gaussian white noise embedded with single (90 ms) or double (210 ms) noise gaps (periods of silence, and the targets of this experiment) was played to both ears (different target sequence in each ear). One ear received the oddball sequence of pure tones (50 ms) at either 500 or 550 Hz (counterbalanced between blocks) (ISI = 450 ms, standard *p* = 0.85 black rectangle, deviant *p* = 0.15 hollow rectangle, respectively). Participants were instructed to pay attention to the targets embedded in the white noise in the left, right or both ears and to ignore the tones. ISI = inter-stimulus interval, L = left ear, R = right ear, Std = standard, Dev = deviant.

### Experimental Design

The twelve experimental trial blocks (T = 3:32min each) were comprised of a total of 380 tones (with deviants always falling within four to 10 standard tones). Participants were instructed to listen for and report target gaps within the white noise stream in either the left channel only, the right channel only, or in either channel (divided attention), and to ignore the tones. Each attention condition was repeated four times and the order of the blocks was pseudo-randomised such that no participant received the same order. When a target was identified in the attended ear(/s) participants responded with a ‘1’ keypress if the gap was singular and a ‘2’ keypress if the gap was doubled. In one third of the blocks oddball tones were played in the attended ear, in another third the tones were played in the ignored ear, and in the remaining third, in which participants divided their attention between ears, the tones were presented to either side, counter-balanced between the left and right across separate blocks. Participants performed all blocks in one testing session of 60 min (42:24 min total task duration plus breaks) with an additional 30 min EEG setup period.

### Task

Participants were seated in front of a computer screen and wore inner-ear buds for the duration of the experiment. Prior to recordings, participants listened to an example auditory stream of 1-min duration, which demonstrated the single and double gaps in the white noise. Each participant then underwent a brief practice session with auditory stimuli consisting of 9 single and 9 double gaps, and a total of 110 tones. Participants were given feedback about their their accuracy in this practice block but not in the experimental blocks. At the beginning of each experimental block, the focus of attention was specified verbally and an arrow (left, right or both directions) remained on the screen for the duration of the block as a reminder. Participants were asked to make their keypresses in response to target gaps as quickly and as accurately as possible, and to ignore any gaps in the uncued ear (in the focused attention condition). Task performance was assessed based on the percentage of correctly detected target gaps and reaction times. Participants with less than 50% overall accuracy (proportion correct) were excluded from further analysis.

### EEG Data acquisition and preprocessing

Continuous EEG data were recorded with a Biosemi Active Two system with 64 Ag/AgCl scalp electrodes arranged according to the international 10-10 system for electrode placement using a nylon head cap. Data were recorded at a sampling rate of 1024Hz. Pre-processing and data analysis were performed with SMP12 (http://www.fil.ion.ucl.ac.uk/spm/). Data were re-referenced to a common reference, down-sampled to 200Hz and high-pass filtered above 0.5 Hz. Eyeblinks were detected and marked using the VEOG channel before the data were epoched offline with a peri-stimulus window of −100 to 400ms. Artefact removal was performed by removing trials marked with an eyeblink and by thresholding all channels at 100uV. Trial data were robustly averaged before being low-pass filtered below 40 Hz and baseline corrected between −100 to 0 ms. We analysed event-related potentials with respect to the onsets of standard and oddball tones, separately for conditions in which the tones were presented in the attended ear, the unattended ear, or in either ear in the divided attention condition.

### Spatio-temporal Image Conversion

Event-related potentials were converted into 3D spatio-temporal volumes per condition and participant. This was achieved by interpolating and dividing the scalp data per time point into a two dimensional 32 × 32 matrix. We obtained one 2D image for every time bin (from 0 to 400 ms in steps of 5ms). These images were then stacked according to their peristimulus temporal order, resulting in a 3D spatio-temporal image volume with dimensions of 32 × 32 × 81 per participant. Data were then smoothed at FWHM 12×12×20 mm^3^.

### Spatio-temporal Statistical Maps

For each participant, the 3D spatio-temporal image volumes were modelled with a mass univariate general linear model (GLM) as implemented in SPM12. We performed between-subject F-contrasts for (1) the main effect of attention, (2) the main effect of prediction and (3) the interaction between attention and prediction. Simple effects were estimated using between-subject t-statistic contrasts. The same statistical analyses were performed on the 3D spatial image volume obtained after source localisation (see below). All sensor effects are reported at a threshold of *p*<0.05 with family-wise error (FWE) correction for multiple comparisons over the whole spatio-temporal volume. For closer inspection of the main effects and interactions obtained at channel Fz (at which predictability effects are typically strongest, Naatanen R and K Alho (1997), we implemented a 1-dimensional GLM approach using SPM12. We restricted our time window from 0 to 400 ms after stimulus onset and, in a separate analysis, between the typical MMN time window of 100 to 250 ms (FWE corrected over the time bins considered).

### Source Reconstruction

We obtained source estimates on the cortical mesh by reconstructing scalp activity with a single-sphere head model, and inverting a forward model with multiple sparse priors (MSP) assumptions for the variance components under group constraints. This allowed for inferences on the most likely cortical regions that generated the sensor-level data. We obtained images from these reconstructions for each of the six conditions in every participant. These images were smoothed at FWHM 12x12x12 mm^3^. We then computed the main effects of attention and prediction, and the interaction (attention × prediction) using conventional SPM analysis. The effect of prediction (t-statistic) is displayed at an uncorrected threshold of *p<*0.001. These weaker significance criteria were used for post-hoc visualisation, once the effects had been established under robust criteria at the scalp level, and we only report regions significant at p<0.05 FWE corrected at the cluster level.

### Statistics

Significance sensor space maps for prediction effects are displayed at p<0.05 corrected for multiple comparisons using family-wise error rate. The interaction map is displayed at p<0.01 uncorrected for the purpose of defining a region of interest for follow up Bayesian Model Selection. Source maps are displayed at p<0.001 uncorrected, but only significant cluster-level p_FWE_<0.05 are reported.

### Bayesian Model Selection

To compare the two models (*Opposition* and *Interaction*; see Introduction) of the effects of attention on prediction (standard and deviant tones) we used the Bayesian Model Selection (BMS) methodology described in Rosa MJ et al. (2010), and adapted here for EEG. For this analysis we discarded trials from the divided attention condition and used only the attended and unattended trials from the focused attention conditions (attend left ear only, attend right ear only) for both standard and deviant tones. We created posterior probability maps (PPMs) from individual participant log-model evidences using a random-effects approach (RFX). Here, the winning model was the one with the highest log-evidence (assuming uniform priors over the models) across participants. We performed this analysis at the sensor and source levels by modelling the data with regressors describing the hypothesised relationships amongst the four different conditions.

Briefly, covariate regressor weights were applied to every participant and trial under the Opposition model, which predicts reductions in ERP amplitudes across conditions in the following order: (1) attended unpredicted, (2) unattended unpredicted/attended predicted and (3) unattended predicted. Next, we specified a second model derived from Kok et al. (2011), the Interaction model, which predicts reductions in ERP amplitudes across conditions in the following order: (1) attended predicted, (2) attended unpredicted, (3) unattended unpredicted and (4) unattended predicted. Voxel-wise whole-brain log-model evidence maps were then created for every participant and model, estimated using the Variational Bayes 1^st^-Level Model Specification methodology described in Penny et al. (2005). Source level maps were further smoothed with a 1 mm half width Gaussian kernel. We used the RFX approach to produce PPMs for both models at the group-level. These maps (displayed at a threshold of probability larger than 75% and 50% for scalp and source, respectively) allowed us to compare which model had the higher probability at each voxel in the brain (and at each time point in the scalp level analysis). Further model comparisons for specific regions at the sensor level were undertaken using brain regions selected a priori from the attention by prediction interaction contrast. At the source, these comparisons were made at the peak coordinates of clusters for each model that exceeded 51%.

### Dynamic Causal Modelling

Source locations were identified based on multiple sparse priors source reconstruction of the overall mismatch (p<0.05 uncorrected threshold). These regions were: bilateral primary auditory cortices (A1; MNI coordinates: left [−42, −24, 34] and right [44, −22, 38]), bilateral inferior temporal gyri (ITG; MNI coordinates: left [−42, −10, −38] and right [44, 0, −42] and left inferior frontal gyrus (LIFG; MNI coordinates: [−50, 32, 0]). First, the basic connectivity architecture was optimised using responses to attended and unattended standards and deviants with no between-trial effects present. This first step considered six competing model structures that differed in the pattern of extrinsic connections and source regions. Next, the pattern of changes in extrinsic connectivity was optimised under this architecture using responses in all four conditions for the Opposition and Interaction models. The family of Opposition models used a between-trial effect of [1, 2, 2, 3] for the attended predicted, predicted attended, unpredicted unattended, and attended unpredicted, respectively. The family of Interaction models, on the other hand used [1, 2, 3, 4] for predicted unattended, unpredicted unattended, unpredicted attended, and attended predicted, respectively. Fifteen competing models were tested, each with a different subset of connections – forward, backward and recurrent – which also included or excluded intrinsic modulations of A1, and a single null model. Finally, the Opposition and Interaction model-dependent changes in intrinsic connectivity were then grouped by families, under the optimised connectivity architecture. In both DCM estimation steps, models were inverted using a 0 to 400 ms peristimulus time window.

## RESULTS

### Behavioural findings on attentional manipulation

Behavioural results for the target detection task – discriminating single- and double-gaps in concurrent white noise streams in each ear – were grouped into unilateral (focused) or bilateral (divided) attention conditions (30 targets over 8 blocks and 60 targets over 4 blocks, respectively). We excluded any participants who did not achieve mean response accuracy greater than 50%. There was no significant difference in response accuracy (*p* = 0.14) between the unilateral (*M* = 71.80%, *SEM* = 5.19%) and the bilateral (*M* = 68.33%, *SEM* = 5.13%) conditions. Participants were significantly faster (*p* = 0.03) to respond in the bilateral (*M* = 748.16 ms, *SEM* = 27.67 ms) than the unilateral conditions (*M* = 779.79 ms, *SEM* = 34.13 ms), likely reflecting a strategy of responding immediately to any target gap when monitoring both ears under divided attention, as opposed to having to select only relevant gaps in the focused attention conditions (filtering out gaps in the ignored ear).

### Attention amplifies prediction errors – single channel analysis

ERPs corresponding to each of the experimental conditions (as well as the MMNs derived from subtracting the standards from the deviants within a condition) were extracted from electrode Fz and compared over time (Figure 3). The N1 and P2 components were plotted as an average across participants and conditions. For this, the lowest time point between 50-150 ms and highest point between 150-250 ms were determined from the omnibus ERP plot (i.e. the mean ERP across all participants and conditions over time). These time indexes +/- 25 ms were then used to find the average ERP per condition. Statistical tests of the N1 components found only a main effect of surprise (F(1,72) = 4.9583, p = 0.0291). Similarly, at P2, there was a main effect of surprise (F(1,72) = 17.5898, p = 7.7001e-05), but no further significant main effects or interactions. In addition, results at Fz from 0 to 400 ms using the 1- dimensional GLM approach revealed a significant main effect of Attention between 290-340 ms (*p*_FWE_cluster_ = 0.006), and a significantly larger prediction error for attended relative to unattended conditions at 115-120 ms (*p*_FWE_cluster_ = 0.020). We then restricted our analysis to the MMN time window (100-250 ms) and again found a significant main effect of Attention but at an earlier period between 200-230 ms (*p*_FWE_cluster_ = 0.028). Moreover, there was a significantly larger prediction error for attended versus unattended conditions between 100-130 ms (*p*_FWE_cluster_ = 0.046). These findings demonstrate that attention amplifies prediction errors.

**Figure 3.**
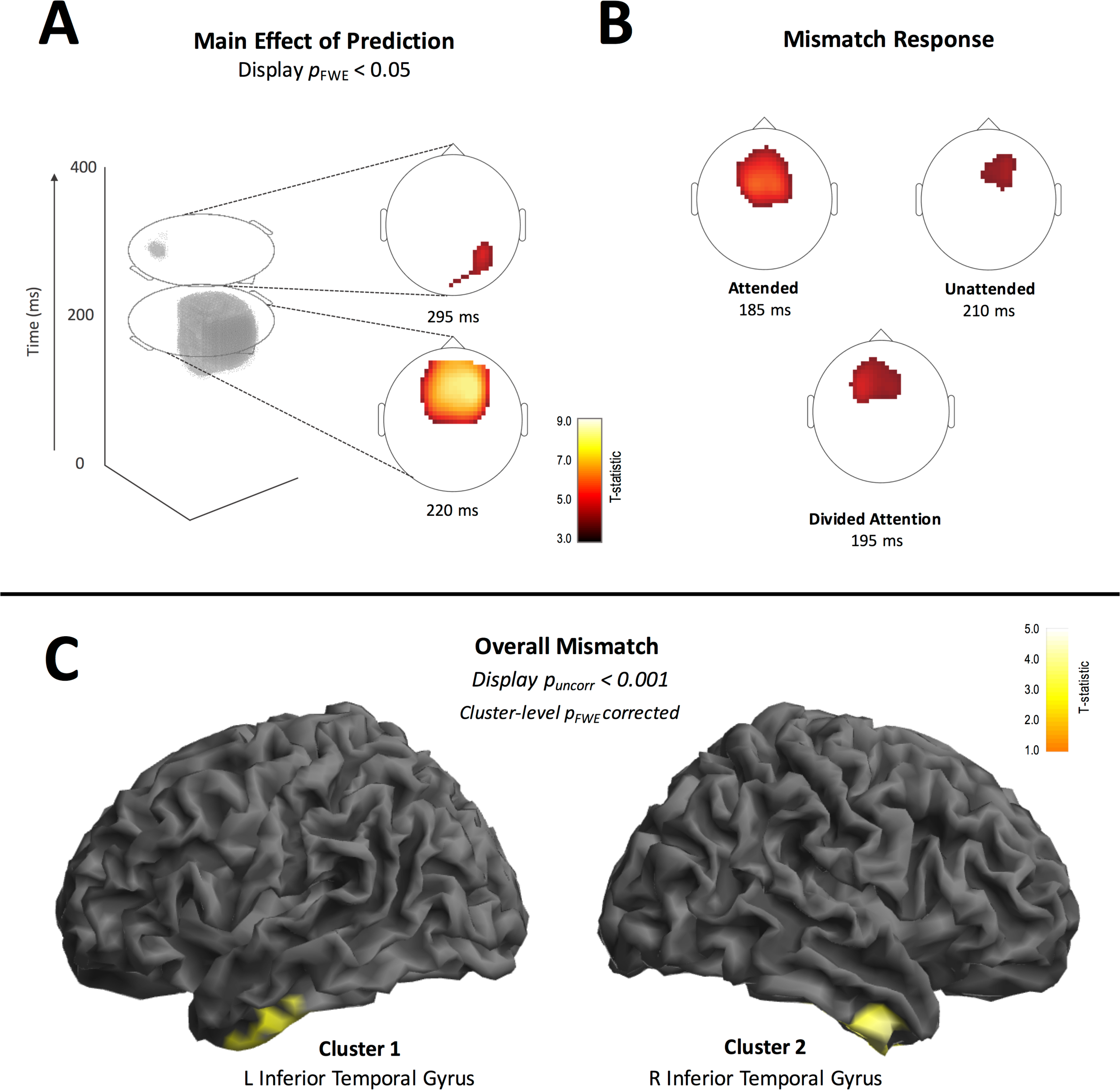
Event-related potentials extracted from electrode Fz for each condition. **(A)** The ERPs for each of the experimental conditions were extracted from electrode Fz and compared over time. The grey shadings indicate the temporal widows during which a significant main effect of attention was found (** corrected for the whole epoch, * corrected within the *a priori* MMN time window). **(B)** ERPs for attended and unattended prediction errors (the MMNs; that is, the difference between unpredicted and predicted) are plotted at electrode Fz. Grey shading indicates the temporal window during which a significan Attention by Prediction interaction was found (* corrected within the *a priori* MMN time window).

### Larger responses to unpredicted than predicted events regardless of attention level – sensor and source space

As shown in Figure 4, the main effect of Prediction, or surprise (standards vs deviants), disclosed several significant components comprised of two late effects. The first late effect was detected from 200-220 ms (peak-level T_max_ = 8.30, cluster-level *p*_*FWE*_ < 0.001; at fronto-central channels). The second late component was observed from 290-295 ms (peak-level T_max_ = 5.02, cluster-level *p*_*FWE*_ = 0.004; at right parieto-occipital channels). We also found simple Prediction effects in all of the attention manipulations, that is, attended (peaking at 185 ms), unattended (peaking at 210 ms), and divided (peaking at 195 ms). While there appeared to be qualitatively differences in the strength and extent of the prediction effects across Attention conditions, the interaction between Attention and Prediction did not survive correction for multiple comparisons.

We then used a multiple sparse priors source reconstruction method to investigate the cortical regions that generated the effects at the scalp level. Statistical parametric maps for source-reconstructed images revealed two significant clusters for the main effect of Prediction in the left ([−42 −10 −38], peak-level T_max_ = 4.14, cluster-level *p*_FWE_ = 0.019) and right inferior temporal gyri ([44 0 −42], peak-level T_max_ = 3.77, cluster-level *p*_FWE_ = 0.023)(Figure 4).

**Figure 4.**
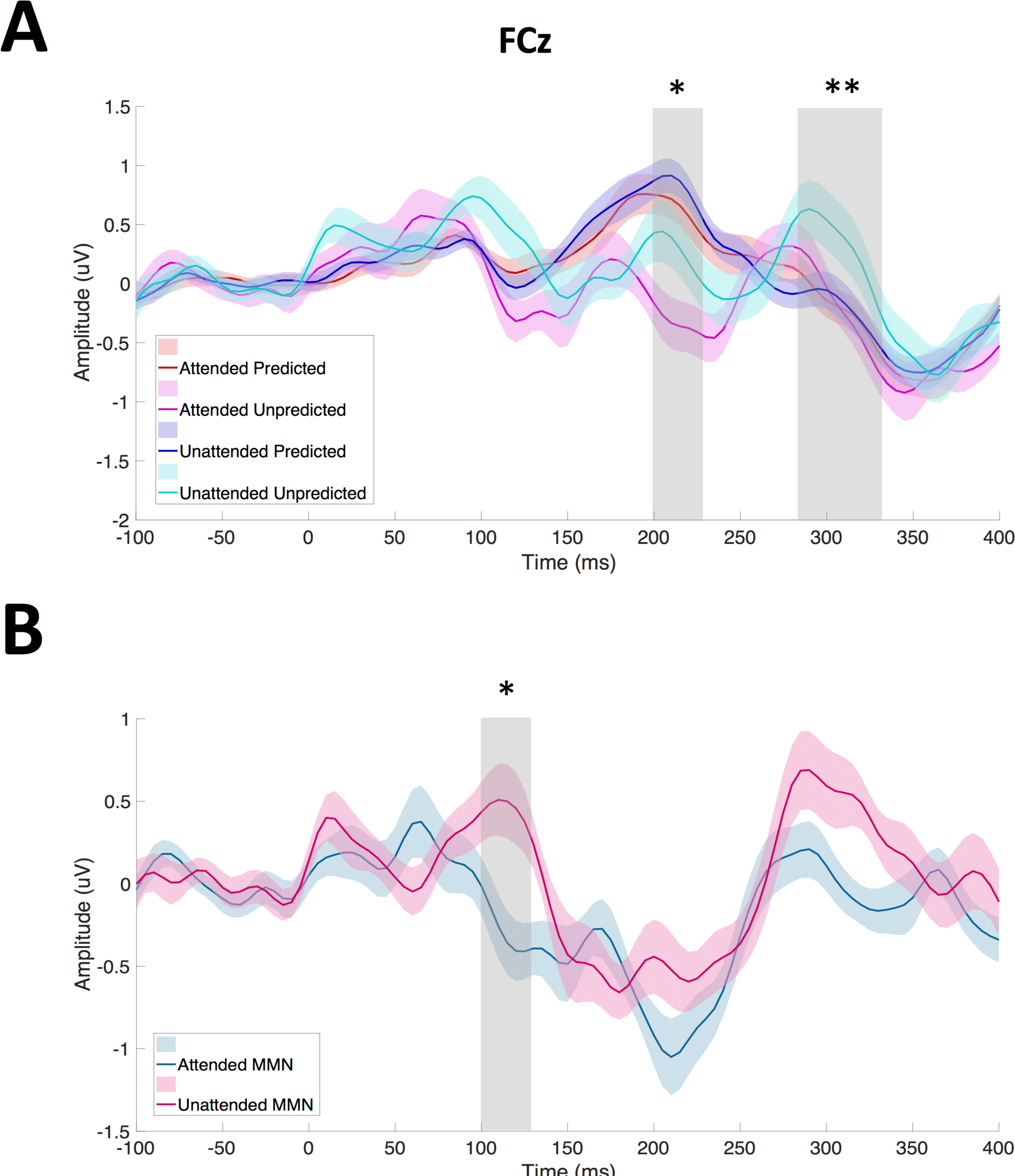
Main and simple effects of prediction at the scalp and source levels. **(A)** Spatio-temporal statistical analysis revealed significant effects of prediction (predicted vs. unpredicted) over fronto-central areas around 220 ms and over posterior parietal areas at 295 ms (displayed at *p*<0.05, FWE whole-volume corrected). **(B)** The effects of prediction across the three attentional manipulations revealed a prediction effect in the attended condition at 185 ms, the divided attention condition at 195 ms, and in the unattended condition at 210 ms, all located fronto-centrally (displayed at *p*<0.05, FWE whole-volume corrected). There was no significant interaction (difference in the MMN between the attention conditions). **(C)** Source reconstruction analysis revealed a main effect of prediction within the left and right inferior temporal gyri. (Displayed at *p*<0.001 uncorrected and FWE corrected at the cluster-level.)

### Opposition wins over Interaction – evidence from posterior probability maps

#### Scalp level

Bayesian Model Selection (BMS) was used to compare the two competing models of the relationship between Attention and Prediction (the *Opposition* or *Interaction* models; see Figure 1). Specifically, we were interested in comparing the strength of neural activation under the different manipulations of attention and prediction. We used random effects BMS to create group-level PPMs for each model, derived from the log-model evidence of each participant, that is, the evidence that a given model (*Opposition* or *Interaction*) generated the data.

As shown in Figure 5, BMS revealed that the ***Opposition*** model (‘*Attention and Prediction oppose*’) was the more likely (>75% model probability) explanation for the data across most fronto-central channel locations at the majority of time points (70 to 210 and 290 to 375 ms). However, the Interaction model (‘*Attention and Prediction interact*’) had a higher probability (>75%) of explaining the data between 170 and 230 ms (i.e. within the MMN time window) at central and lateral parietal channel locations. Thus, the relationship between Attention and Prediction differed depending on both the time point and scalp location; although more often than not, Attention and Prediction had opposing effects.

**Figure 5.**
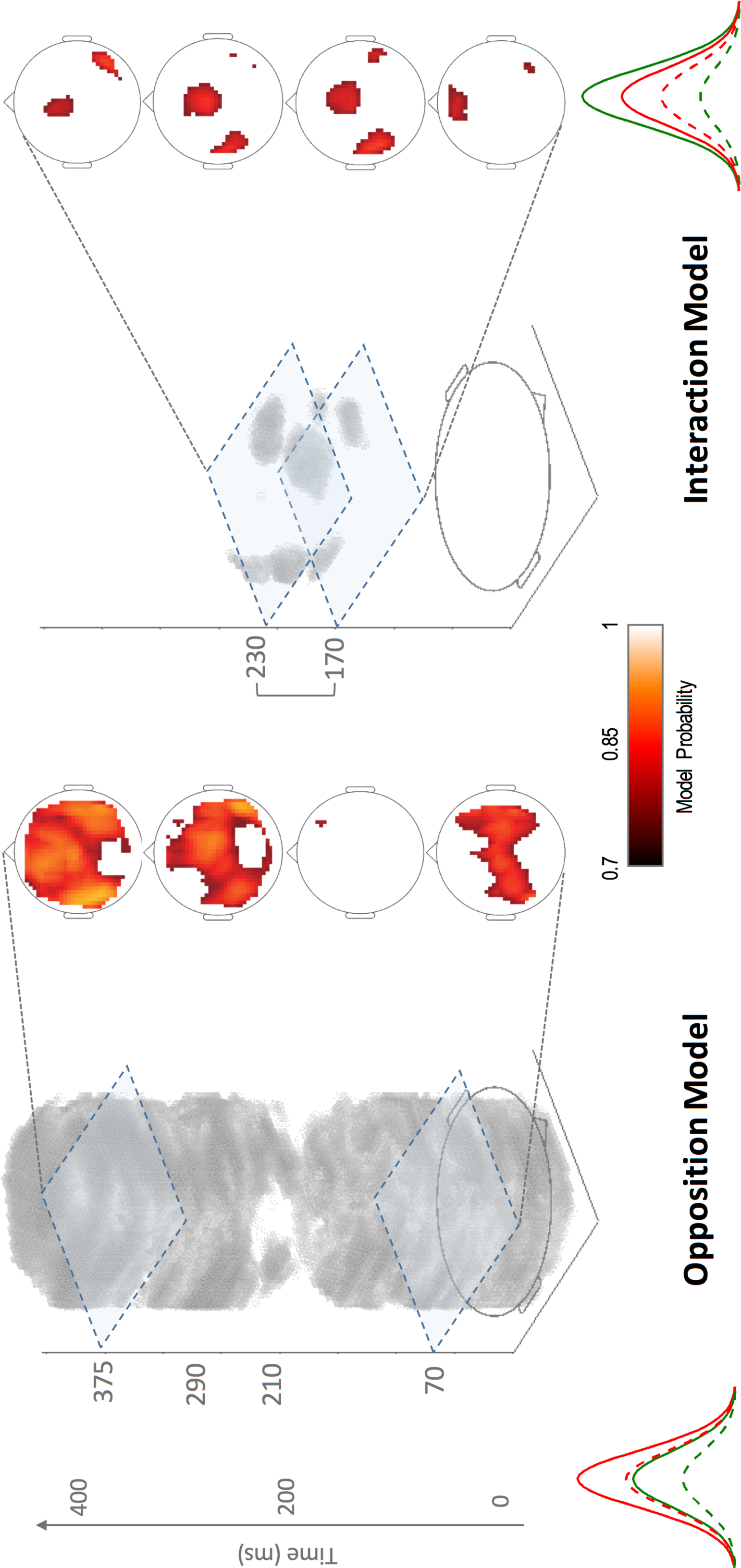
Scalp Posterior Probability Maps of the Opposition and Interaction models over space and time. Maps display the posterior probability for both models, thresholded at probability >75% over space and time. Scalp maps show the four timepoints with the largest significant clusters. The *Opposition* model wins (*Attention and Prediction oppose*) across most fronto-central channels at the majority of time points (70-210 and 290-375 ms). The *Interaction* model wins (*Attention and Prediction interact)* at the fronto-central and lateral parietal regions of the scalp (channel locations) between 170-230 ms.

The fact that the Interaction model won within the MMN window and yet we did not find a significant interaction in the classic GLM analysis could perhaps be explained by a Prediction by Attention interaction effect that did not survive correction for multiple corrections. We further examined a potential interaction effect, hindered perhaps by a rather conservative multiple comparison correction procedure. Firstly, we used more lenient, uncorrected peak-level statistics to select two small interaction clusters at 175 ms (peak-level F_max_ = 5.79, peak-level *p*_*uncorr*_ = 0.004; at central channels) and 360 ms (peak-level F_max_ = 5.45, peak-level *p*_*uncorr*_ = 0.006; at right parietal channels – see Figure 6). We then took the spatio-temporal coordinates of these clusters and extracted the posterior probability of each model at that particular location. We constructed a 10^3^ cube around these coordinates and took the average posterior probability of each model over that volume. Our reasoning was that if an interaction between Attention and Prediction were present in the data, then the *Interaction* model would have a higher posterior probability compared with the *Opposition* model at these coordinates. We found that at 175 ms over fronto-central channels there was a negligible difference between the *Opposition* and *Interaction* models, with 48% and 52%, respectively (Figure 6). However, at 360 ms over the right lateral parietal area, the *Opposition* model probability far exceeded that of the *Interaction* model, with a value of 80%. Thus, Attention and Prediction appear to have opposing effects later in time.

**Figure 6.**
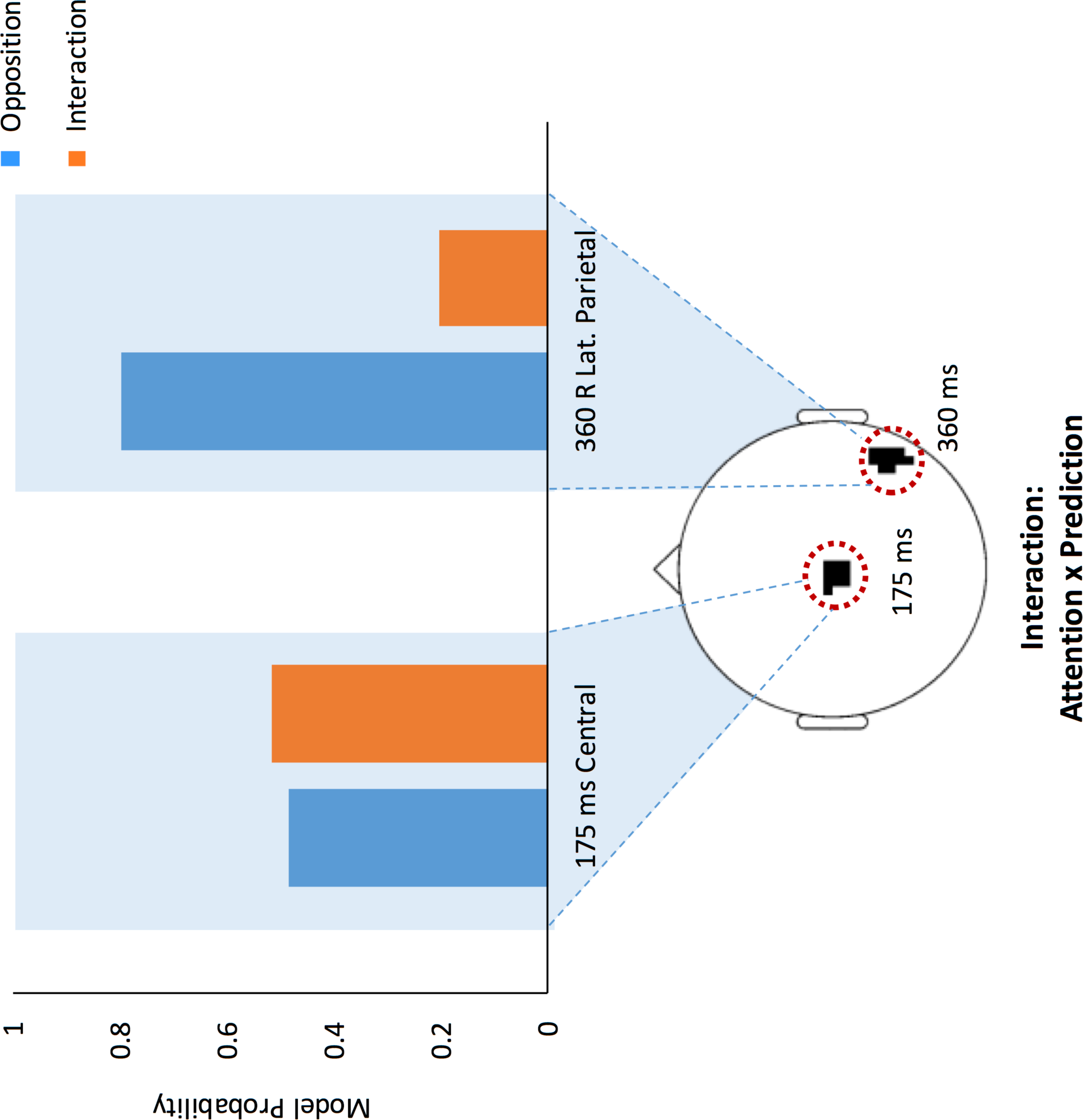
Bayesian Model Comparison within the spatio-temporal clusters extracted from the Prediction by Attention interaction. We extracted model probabilities using the coordinates (scalp location and time points) of two clusters from the Interaction results (based on the liberal threshold of p<0.001 uncorrected). If interaction effects were present, the *Interaction* model would be more likely to win over the *Opposition* model at these coordinates. At 175 ms (within the MMN time window) and over central electrodes, there was a very slight advantage for the *Interaction* over the *Opposition* model. However, at 360 ms over right lateral parietal channels, the *Opposition* model probability far exceeded that of the *Interaction* model, with a probability of 80%.

#### Source level

Finally, we applied the same BMS technique employed at the sensor level to our source reconstructed results. BMS revealed that the *Opposition* model had the higher model probability and larger clusters at the source (Figure 7). The *Opposition* model achieved >50% model probability in the left middle temporal gyrus (cluster size; K_E_ = 82) and right inferior temporal gyrus (cluster size; K_E_ = 288). Conversely, the *Interaction* model achieved >50% model probability in a smaller cluster in the left middle temporal gyrus (cluster size; K_E_ = 32). We then compared the model probabilities at the centre of these clusters and showed that the *Opposition* model was more probable than the *Interaction* model in the left middle temporal and right inferior temporal gyri (winning with 82% and 78% probability, respectively). Furthermore, model probabilities extracted from the peak of the *Interaction* model cluster showed a slight advantage for the *Interaction* over the *Opposition* model (with 57% probability for the *Interaction* model).

**Figure 7.**
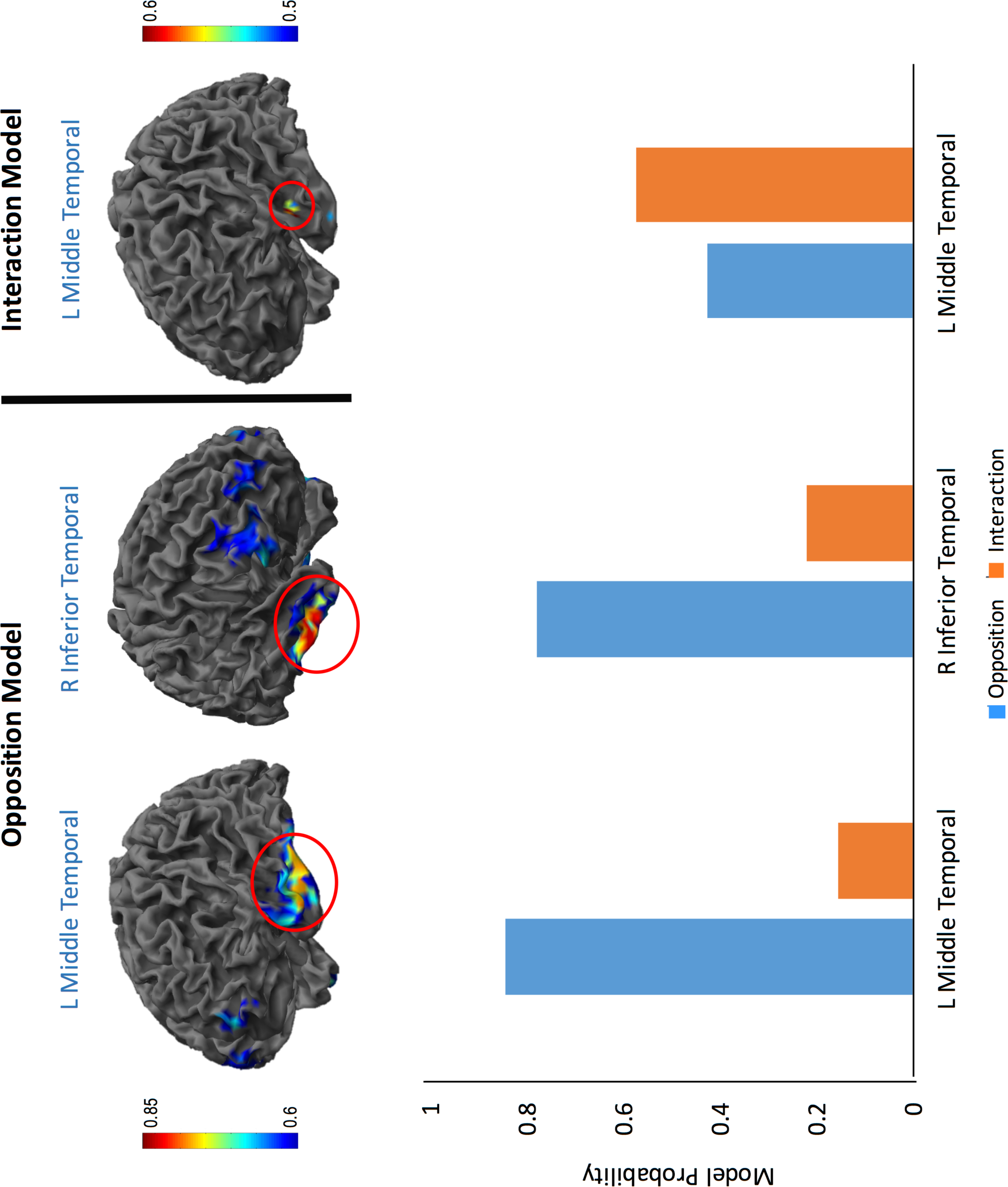
Source Posterior Probability Maps of the Opposition and Interaction models (top) and model probabilities for the three major clusters of the two models (bottom). BMS was used for model inference at the group-level at the source. Here, the *Opposition* model achieved >50% model probability in the left middle temporal (cluster size; K_E_ = 82) and right inferior temporal (cluster size; K_E_ = 288) gyri. The *Interaction* model achieved >50% in a small cluster in the left middle temporal gyrus (cluster size; K_E_ = 32). Overall, the *Opposition* model achieved higher probability over a larger number of voxels. Extraction of model probabilities from the peak of the *Opposition* clusters showed that this model won with 82% probability in the left middle temporal gyrus and with 78% probability in the right inferior temporal gyrus. Model probabilities extracted from the peak of the *Interaction* cluster showed there were minimal differences between either model at this location, with 57% posterior probability for the *Interaction* model. Note the differences in the colour map scales between the *Opposition* and *Interaction* models.

#### Dynamic Causal Modelling

The prior location of the cortical sources included in our DCMs was based on MSP source reconstruction of ERPs corresponding to the 4 conditions (attended standards, attended deviants, unattended standards, and unattended deviants) of the Overall Mismatch. Statistical parametric maps were inspected at a more liberal threshold of p<0.05 (uncorrected) to identify candidate neural sources of the effects observed on ERP amplitude (Auksztulewicz R and K Friston 2015). Following the selection of candidate sources, model structure was optimized by comparing 6 alternative connectivity models using data from each of the experimental conditions, with no between trial effects present (i.e. 1 to 3 levels of connectivity, with or without lateral connections between the inferior temporal gyri). Results indicated that the best model included recurrent connection amongst all regions, that is, inputs to LA1 and RA1, with LA1 connected to LITG, and LITG connected to LIFG, as well as connections linking RA1 and RITG, and lateral connections between LITG and RITG. The selected model was then used to further optimise condition specific changes in the extrinsic connectivity by comparing the types of extrinsic connections present. Here, 15 alternative models were compared, with each condition-specific model (*Opposition* and *Interaction*) modulating a different subset of extrinsic connections (forward only, backward only or forward and backward). A null model with no modulations on any connection was also tested. These models were fitted to each participant’s data to explain observed differences in ERP amplitude. Random-effects Bayesian model selection revealed that the *Opposition*-model with modulation of forward connections outperformed all other models.

## DISCUSSION

In this study, we adjudicated between two alternative computational models of the effect that spatial attention has on expectations. Using Bayesian model comparison of scalp posterior probability maps we found that, except for an early time window (within the typical MMN), the *Opposition* model won over the *Interaction* model. This suggests that, for the most part, attention provides an equivalent boost to neuronal responses to predicted and unpredicted stimuli. Similarly, at the source level we found stronger evidence for the *Opposition* model underlying a fronto-temporal network. We investigated this further with DCMs that employed trial-dependent plastic changes according to either the *Opposition* or the *Interaction* model. In agreement with the model-based scalp and source analysis, we found that the family of *Opposition* models better explained the data. Classic SPM analysis of spatio-temporal maps revealed an effect of prediction across and within all attentional manipulations, which peaked within the typical MMN time window and at fronto-central channels. This effect was statistically greater in the attended compared with the unattended conditions at the single channel level, where MMN is typically seen, suggesting that attention amplifies prediction errors. At the whole spatiotemporal map level, however, this interaction effect did not survive correction for multiple comparisons over the whole space-time, despite the appearance of somewhat larger clusters for the attended than the unattended condition,

Our finding of a prediction error effect in all attention conditions (attended, unattended and divided) is in agreement with a vast body of work suggesting that the MMN is elicited regardless of attention, and hence is ‘pre-attentive’ in nature (Naatanen R *et al*. 2001). This is in contradistinction to Auksztulewicz R and K Friston (2015), who did not find an effect of prediction in the absence of attention (although this might have been due to a lack of power, as very few trials were included). Again, our finding of a prediction error effect regarless of attention is opposite to Todorovic A et al. (2015), who found that while beta synchrony decreased with expectation in the unattended condition, no difference was found in the attended condition. The latter is seemingly at odds with the idea that attention amplifies prediction errors as previously shown (Jiang JF et al. 2013; Auksztulewicz R and K Friston 2015), and as revealed in the current study. A number of factors could be explain such conflicting results. Perhaps most importantly, very different paradigms and measures were employed across the relevant experiments. Both our study and that of Auksztulewicz R and K Friston (2015) investigated the effects of attention and prediction on evoked responses in an oddball paradigm, whereas Todorovic A *et al*. (2015) focused on endogenous oscillatory activity. Moreover, both Auksztulewicz R and K Friston (2015) and Todorovic A *et al*. (2015) manipulated temporal attention, whereas here we manipulated spatial attention. Moreover, in our experiment attention and prediction were manipulated within the same spatial location (left or right ears), but were drawn toward independent auditory ‘objects’ (noise for the attention task, and tones for the concurrent oddball stream). By contrast, the aforementioned studies (and that of Kok P *et al*. (2012)) manipulated attention and prediction within the same (visual or auditory) object. Future work should test whether manipulating attention and prediction for common versus independent stimuli alters the extent to which they interact.

In this work we directly compared two competing models of the effects of attention on expectations – the *Interaction* and *Opposition* models – put forward in Kok P *et al*. (2012). The data in that study were consistent with the Interaction model when considering regions of the visual cortex (V1, V2 and V3). Here, however, we took a different approach by implementing the models computationally and directly testing them against our data. By using Bayesian model comparison of statistical maps of EEG activity, and DCMs for ERPs, we were able to quantify how likely each of these two models was at every point of space and time at the scalp level, at each voxel in source space, an in the trial-dependent plastic changes within a cortical network. The Opposition model was unambiguously favoured in our data at every level, i.e., scalp, source, and network. At the network level we found that the plastic changes according to the *Opposition* model were more pronounced in forward connections. This is consistent with the idea that attention boosts, or heavily weights, prediction errors, which are then conveyed upward in the cortical hierarchy. Such prediction errors signal the need to update an internal perceptual model of the world, in turn prompting learning. While these findings are mostly at odds with those in Hsu YF *et al*. (2014) and Kok P *et al*. (2012), there was a narrow window of agreement in which the *Interaction* model was better at explaining the data at the scalp level, perhaps tellingly within the MMN time frame. This is an interesting finding, as it seems to point to a tonic *Opposition* effect between Attention and Prediction, and a phasic *Interaction* effect. Again, there are differences in both the type of paradigm and the neuronal measures between our study, which used EEG, and the experiment of Kok P *et al*. (2012), which used fMRI. Although Attention was manipulated spatially in both studies, in our study it was directed towards a different (instead of the same) object. Moreover, our Prediction manipulation was learnt from the sequence of stimuli, rather than instructed (as in Kok P *et al*. (2012).

In conclusion, our findings provide empirical evidence for a computational model of the opposing interplay of attention and expectations in the brain. These opposing effects are manifested in neuronal activity and in plastic changes within a fronto-temporal network engaged in sensory prediction errors. We demonstrate that attention boosts neuronal responses to predicted and unpredicted stimuli, and replicate the finding that attention boosts prediction errors, in keeping with the predictive coding framework (Rao RP and DH Ballard 1999; Friston K 2005). Finally, we demonstrate that prediction errors are elicited regardless of one’s state of attention, providing further support to the idea of a pre-attentive nature of change detection systems in the brain (Naatanen R *et al*. 2001).

**Figure 8.**
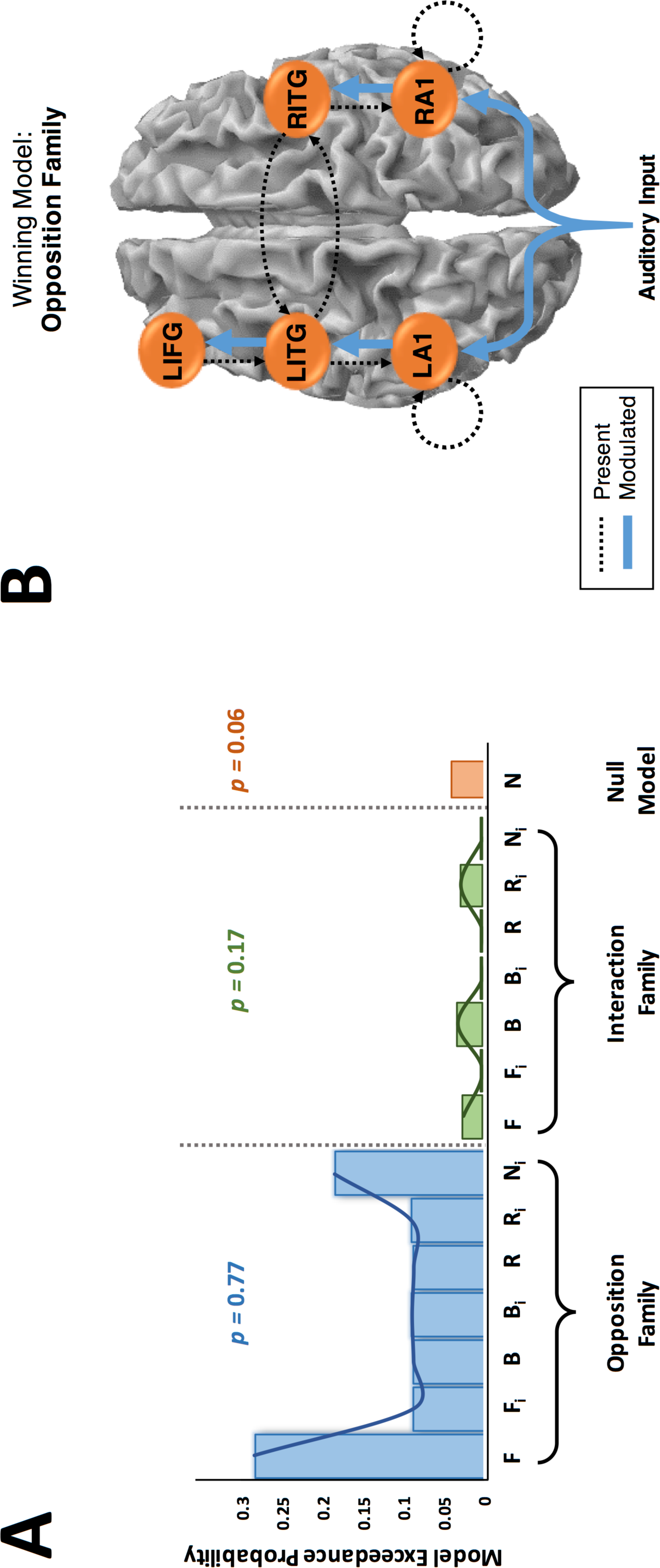
Dynamic Causal Modelling hypotheses testing for plastic changes according to the Opposition and Interaction families of models. Source locations were identified as bilateral primary auditory cortices (LA1 and RA1), bilateral inferior temporal gyri (LITG and RITG) and left inferior frontal gyrus (LIFG). **(A)** Extrinsic connectivity was optimised using responses in all four conditions under the *Opposition* or *Interaction* model. Fifteen competing models were tested, each with a different subset of trial-specific modulation of connections, according to the *Opposition* and *Interaction* models on forward (F), backward (B) and recurrent (R) connections (with and without intrinsic modulations of A1 (subscript i)), as well as a null model precluding any modulations. Summed model exceedance probabilities across each family show the winning family as the *Opposition* Family (left; blue). **(B)** The winning model architecture had recurrent connections between all regions, intrinsic modulation of A1, lateral connections between bilateral ITG, and included trial-dependent modulations according to the *Opposition* model in the forward connections (blue lines).

## Acknowledgements

This work was funded by an Australian Research Council (ARC) Discovery Early Career Researcher Award (DE130101393) and a University of Queensland Fellowship (2016000071) to MIG, an ARC Australian Laureate Fellowship (FL110100103) to JBM, the ARC Centre of Excellence for Integrative Brain Function (ARC Centre Grant CE140100007) to MIG and JBM, and an ARC Special Research Initiative - Science of Learning Research Centre (SR120300015) to JBM. We thank the volunteers for participating in this study and Maria Joao Rosa for discussions.

